# randPedPCA: Rapid approximation of principal components from large pedigrees

**DOI:** 10.1101/2025.03.17.643624

**Authors:** Hanbin Lee, Rosalind F. Craddock, Gregor Gorjanc, Hannes Becher

**Affiliations:** Department of Statistics, University of Michigan, Ann Arbor, MI, USA; The Roslin Institute, Royal (Dick) School of Veterinary Science, University of Edinburgh, Midlothian EH25 9RG, UK

## Abstract

**Background:** Pedigrees continue to be extremely important in agriculture and conservation genetics with the pedigrees of modern breeding programmes easily comprising millions of records. The structure of such pedigrees is challenging to visualise. Being directed acyclic graphs, pedigrees can be represented as matrices. Common choices are the numerator relationship matrix, ***A***, and the adjacency matrix ***T***. With these matrices, the structure of pedigrees can then, in principle, be visualised via principal component analysis (PCA). However, the naive PCA of matrices for large pedigrees is challenging due to computational and memory constraints.

**Results:** We present the open-access R package `randPedPCA` for rapid pedigree PCA using sparse matrices and randomised linear algebra. Our rapid pedigree PCA builds on the fact that matrix-vector multiplications with the numerator relationship matrix can be carried out implicitly using the extremely sparse inverse Cholesky factor of the numerator relationship matrix. We demonstrate the utility of randPedPCA using several simulated datasets of pedigrees and SNPs. We then demonstrate the performance of `randPedPCA` by analysing the pedigree of the UK Labrador Retriever breeding population of almost 1.5 million individuals.

**Conclusions:** The structure of pedigrees can be efficiently and rapidly visualised using scatter plots of principal component scores. For large pedigrees, this is considerably faster than rendering plots of a pedigree graph.

## 1 Background

Pedigrees continue to be extremely important in the fields of selective breeding and conservation genetics. Because pedigrees indicate the expected genetic similarity between individuals, they can be used to estimate the degree of inbreeding and coancestry, and the expected genetic merit of individuals when coupled with phenotype data. The pedigrees of long-established breeding programmes can easily contain millions of records. This sheer scale makes them difficult to visualise. To aid visualisation of large pedigrees from selective breeding programmes, Garbe and Da (2003) plotted a pedigree graph and collapsed large progeny groups into nodes, which can be very effective in some, but not all populations or species.

Pedigrees are directed acyclic graphs, which can be represented as a matrix. The most natural representation is perhaps the sparse adjacency matrix (e.g. Kepner and Gilbert, 2011). Alternatively, to encode the implied genetic relations between individuals, one can use the sparse adjacency matrix and, given a statistical model, derive the corresponding dense covariance (numerator relationship) matrix (e.g. Lauritzen, 1996). One popular tool for visualising matrices (and multidimensional data in general) is principal component analysis (PCA) (Hotelling, 1933; Pearson, 1901). Technically, PCA is based on an eigen decomposition of a covariance matrix or a singular value decomposition (SVD) of the underlying data matrix (Golub and Van Loan, 2013; Jolliffe, 2002). For a symmetric data matrix, these two decompositions are closely related, as the eigenvectors and singular vectors coincide, and the singular values correspond to the absolute values of the eigenvalues. PCA produces a new matrix, whose columns are orthogonal linear combinations of columns in the data matrix (Golub and Van Loan, 2013; Jolliffe, 2002). The columns are referred to as principal component scores, or simply principal components (PCs). The first principal component accounts for the largest amount of variance shared across the columns of the data matrix, and the subsequent principal components account for increasingly smaller amounts. For many datasets, it is sufficient to obtain the first few principal components to provide an overview of the structure of the data matrix.

Because pedigrees can be represented as matrices, they can also be visualised with PCA. However, few examples of pedigree PCA have been published (Honda et al., 2002; Sneller, 1994; Souza and Sorrells, 1989), with a conspicuous absence of applications to large pedigrees. State-of-the-art genetics research is now based on highly informative genome-wide marker genotypes or whole-genome sequence data (Hanotte et al., 2002; Menozzi et al., 1978; Novembre and Stephens, 2008; Patterson et al., 2006), with theory for understanding PCA in terms of genetic ancestry (McVean, 2009; Peter, 2022; Zheng and Weir, 2016). However, we still have large pedigrees, some with millions of individuals, which we would like to visualise efficiently. There are two key reasons for the absence of PCA for large pedigrees. First, the naïve encoding of a pedigree as an additive relationship matrix (a dense covariance matrix) incurs quadratic storage complexity. Second, decomposing large dense matrices also has a significant compute complexity. For example, the SVD of a dense matrix has *O* (*min* (*nk*^2^, *n*^2^*k*)) complexity for an *n* × *k* matrix, while the truncated SVD of a dense matrix has *O*(*nkr*) complexity to obtain the first *r* components (Eckart and Young, 1936). There are also SVD algorithms that can operate on sparse matrices with *O*(*zk*) compute complexity for *z* non-zero elements in the matrix (Larsen, 1998; Lehoucq et al., 1998).

Henderson (1975, 1976) and Quaas (1976) developed an efficient method to compute the Cholesky factor of the dense additive relationship matrix, and of its sparse inverse, the precision matrix. The Cholesky factor of the precision matrix is a function of a part of the sparse pedigree adjacency matrix and a vector of individuals’ Mendelian sampling variance, in line with the underlying quantitative genetic model (Kennedy et al., 1988; Quaas, 1988; Thompson, 1979; Wright, 1921, 1922). The number of non-zero elements in the precision matrix and its Cholesky factor are therefore proportional to the number of individuals, enabling efficient linear algebra operations with large pedigrees (Aguilar et al., 2011; Colleau, 2002; García-Cortès et al., 2010; Gengler et al., 2007; Henderson, 1976; Meuwissen and Luo, 1992; Sargolzaei et al., 2005; Strandèn and Mäntysaari, 2020; Thompson, 1979; Tier, 1999).

While efficient linear algebra operations on pedigrees are possible, we are not aware of an algorithm for a scaleable PCA to visualise and study large pedigrees. Recent advances in randomised numerical linear algebra have led to the development of efficient algorithms for common matrix operations (Murray et al., 2023). For example, randomised SVD can rapidly approximate truncated SVD of a dense or sparse matrix, including matrices defined implicitly by matrix-vector multiplication (Halko et al., 2011; Voronin and Martinsson, 2016). This randomised algorithm approximates the column space of a matrix using random test vectors and then computes the SVD on this low-dimensional approximation to obtain the dominant singular vectors and singular values. Despite the randomness of the algorithm, it behaves almost deterministically with high efficiency for matrices with rapidly decaying eigenvalues. Critically, the algorithm only implicitly accesses the original matrix via matrix-vector products, without requiring the full matrix.

The aim of this contribution is to implement an efficient algorithm for rapid PCA of large pedigrees by using randomised SVD and the sparse Cholesky factor of the pedigree precision matrix. Specifically, we implicitly represent the core matrix-vector product in the randomised SVD by solving a triangular system of equations based on the sparse pedigree precision Cholesky factor and the vector. The number of operations required for this operation is proportional to the number of individuals in the pedigree, hence scaling to millions of individuals. Another advantage is that the resulting algorithm has a low memory requirement and does not suffer from slow disk access, thus bypassing the significant computational bottlenecks faced by genotype PCAs. We benchmarked the algorithm with simulated data and a large empirical pedigree, and implemented it in the freely available `randPedPCA` R package. Taken together, this work complements the existing toolbox for analysing large pedigrees.

## 2 Implementation

The central piece of our rapid pedigree PCA is an approximate decomposition of the pedigree relationship matrix ***A***, obtained via randomised SVD (Halko et al., 2011). This randomised SVD operates on the sparse Cholesky factor, ***L***^*−*1^, of the inverse of the relationship matrix ***A***^−1^ (Henderson, 1975, 1976; Quaas, 1976). Following the pedigree quantitative genetic model (Kennedy et al., 1988; Quaas, 1988; Thompson, 1979; Wright, 1921, 1922), ***A*** = ***LL***^*T*^ = ***T DT*** ^*T*^ is a covariance (coefficient) matrix, ***L*** is the Cholesky factor of ***A, T*** ^−1^ is closely related to the pedigree graph adjacency matrix, and ***D*** is a diagonal matrix of Mendelian sampling variances. Throughout this work, we refer to the additive relationship matrix, which is also known as the numerator relationship matrix, as the relationship matrix or covariance matrix. We approximate the decomposition of ***A*** with a randomised SVD, which approximates the ‘structure’ of ***A*** by creating an orthonormal range matrix, ***Q***, with a reduced number of columns (Halko et al., 2011). However, we do not operate directly on ***A***. Instead, we use the Cholesky factor of its inverse, ***L***^*−*1^, which is sparse and triangular. The number of non-zero entries in ***L***^−1^ is linear with the size of the pedigree and the matrix can be computed directly from a pedigree (Henderson, 1975, 1976; Quaas, 1976), for example, using the `pedigreeTools` R package (Vazquez et al., 2024). In the following, we describe the key algorithms that enable the rapid PCA of large pedigrees. These are: (i) indirect matrix-vector multiplication of ***A*** with a vector, (ii) centring the implicit data matrix, (iii) randomised SVD for pedigree PCA, and (iv) trace estimation for total variance.

To efficiently multiply ***A*** with a vector ***x***, we exploit the sparsity of ***L***^*−*1^. This is based on a wellknown result (Colleau, 2002), which we describe here for completeness. Note that ***Ax*** = (***LL***^*T*^) ***x*** =***L*** (***L***^*T*^ ***x***) = ***b***. The multiplication on the right, 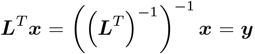, is a backward substitution with (***L***^*−*1^) ^*T*^ in the system of equations (***L***^*−*1^) ^*T*^ ***y*** = ***x***, because (***L***^*−*1^) ^*T*^ is an upper triangular matrix. The remaining multiplication, ***Ly*** = (***L***^*−*1^) ^−1^ ***y*** = ***b***, is a forward substitution with ***L***^−1^ in the system of equations ***L***^*−*1^***b*** = ***y***. Both substitutions cost *O*(*n*) operations and are available in standard linear algebra libraries (Anderson et al., 1999; Golub and Van Loan, 2013). For example, the `spam` R package (Furrer, Flury, et al., 2022; Furrer and Sain, 2010) has efficient implementations of these operations for sparse matrices. We summarise this routine in Algorithm 1. In the context of randomised SVD, ***x*** is a random vector. Instead of performing the matrix-vector multiplication repeatedly with different random vectors ***x***, we carry out one pass with the test matrix **Ω**, whose columns give different vectors ***x***.

### Algorithm 1

Efficiently multiplying the relationship matrix ***A*** with a vector ***x*** via the Cholesky factor, ***L***^*−*1^, of the precision matrix

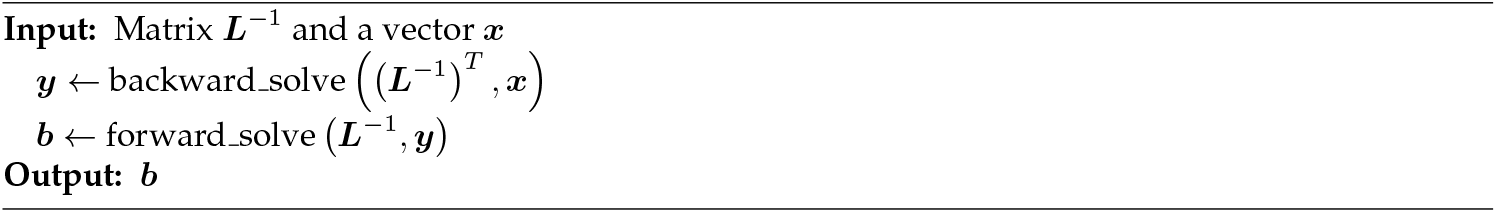

Now, we show how to efficiently ‘centre’ the pedigree relationship matrix ***A*** for PCA. ***A*** is closely related to the genotype relationship matrix ***G***, which is commonly used for PCA in genetics. The genotype relationship matrix is commonly calculated as 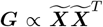, where ***X*** is a genotype (data) matrix with loci in columns and individuals in rows and 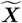 is the column-centred genotype matrix (VanRaden, 2008). Because ***G*** is computed from a centred genotype matrix, we refer to it as ‘centred’, but note that its column means are non-zero. The genotype *x*_*i,l*_ of individual *i* at locus *l* can be viewed as a random variable with respect to the meiotic process on a pedigree. For a given infinite pedigree, the expectation of ***G*** according to the meiotic process is the ‘centred’ ***A***:

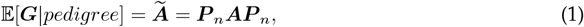

where *n* is the number of individuals, 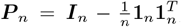, and **1**_*n*_ is a column vector of *n* ones. Hence, the PCAs of ***G*** and 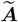 are closely connected, which we demonstrate in the Results section. Using the right-hand side of equation (1), we can efficient and implicitly ‘centre’ ***A***. Consider 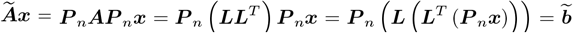. The right-hand multiplication of ***P*** _*n*_ to a vector ***x*** is simply subtracting the mean of the elements from the vector, that is, centring the vector, so we do not need to explicitly form ***P*** _*n*_. This gives us Algorithm 2 for efficiently multiplying the ‘centred’ relationship matrix 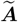 with a vector ***x***.

### Algorithm 2

Efficiently multiplying the ‘centred’’ relationship matrix ***A*** with a vector ***x*** via the Cholesky factor, ***L***^*−*1^, of the precision matrix

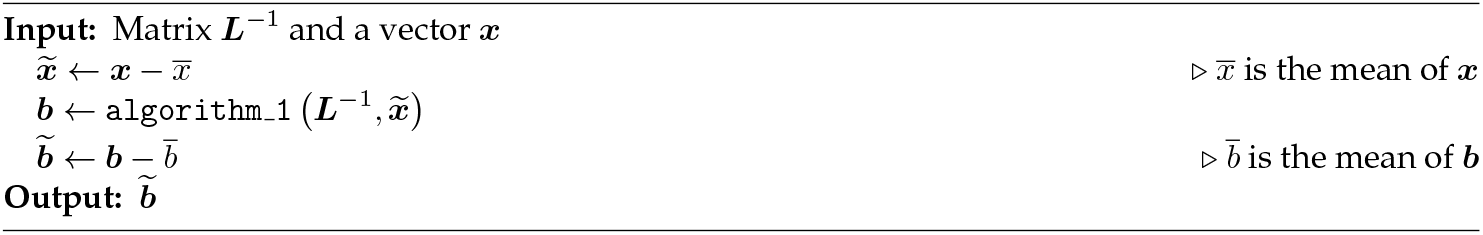

With the essential matrix-vector multiplication algorithms in hand, we can establish the randomised SVD algorithm for rapid pedigree PCA following Halko et al. (2011). The algorithm multiplies ***A*** to a random matrix of independent Gaussian variables **Ω** ∈ ℝ^*n×l*^ via Algorithm 1, and applies QR decomposition (with *O* (*nl*^2^) compute complexity) to obtain an approximate rank-*l* or-thogonal range matrix ***Q*** ∈ ℝ^*n×l*^. To increase accuracy, the integer *l* is chosen to be larger than the desired number of principal components, *k*. According to Halko et al. (2011), there is no need to set *l* to more than 2*k*. This multiplication step can be repeated multiple times to improve the range matrix, where subsequent steps multiply ***A*** to ***Q*** from the previous step instead of **Ω**. Once ***Q*** is obtained, we compute ***B*** = ***Q***^*T*^ ***A***, see Halko et al. (2011), efficiently via backward substitution with (***L***^*−*1^) ^*T*^. Given that the number of desired columns, *k*, in ***B*** is ≪ *n*, it is then possible to run an ordinary SVD, which returns: a *n×l* matrix of left singular vectors, ***U***, a*×l* 1 vector of singular values, ***d***, and an *l×l* matrix of the right singular vectors, ***V*** . From each of these, we then use only the *k* first columns (or elements). The matrix-vector product ***QUd***^2^ gives the *k* approximate principal component scores of ***A*** (Golub and Van Loan, 2013; Jolliffe, 2002). While fully customisable, we have set default values of *k* = 10 and *l* = 15 in our implementation, which we found to work well on simulated and empirical data. We square the singular values because the entries of ***d*** are the singular values of ***L*** (due to using the backward substitution to compute ***B***). The vector ***d*** also represents the standard deviation of each approximated principal component. This randomised SVD is summarised in Algorithm 3.

### Algorithm 3

Approximate PCA of relationship matrix ***A*** (possibly ‘centred’) via the randomised SVD of the precision matrix’s Cholesky factor ***L***^*−*1^

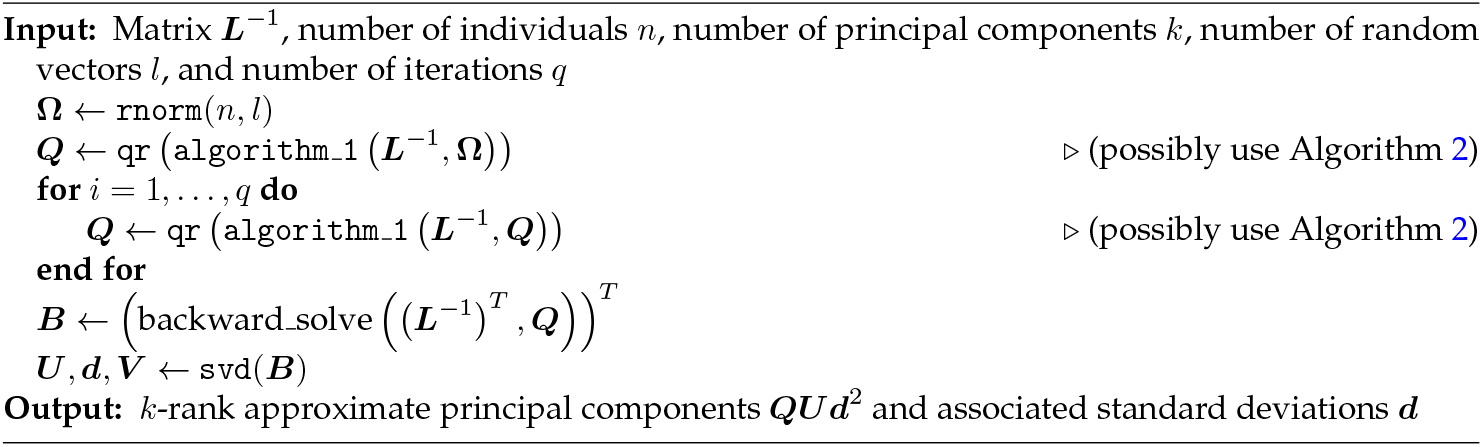

PCA implementations usually return the principal component scores and associated standard deviations, which enables the reporting of the proportion of total variance captured by each principal component. The total variance is the sum of the squared standard deviations of the principal components, or equivalently the trace of covariance matrix. Because truncated SVD returns only a subset of the principal components, and associated standard deviations, the total variance and variance proportions cannot be computed. To calculate the total variance, we use the Meuwissen and Luo (1992) algorithm, implemented in the `pedigreeTools` R package (Vazquez et al., 2024). This computes all the individuals’ inbreeding coefficients. We then sum these and add *n* to the sum, all without forming the matrix ***A***. When using the ‘centred’ relationship matrix 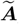, we efficiently compute its trace as 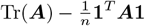. We derived this expression by applying the trace operator to equation (1):

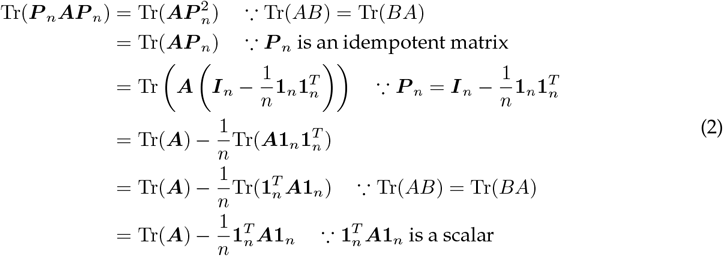

The trace of the ‘centred’ relationship matrix thus depends on the trace of the non-centred version, which is straightforward to compute from the pedigree. If the user only has access to the ***L***^−1^ matrix, and not the pedigree, the trace of ***A*** must be estimated. Inspired by the randomised SVD algorithm, we initially tested the popular Hutchinson (1990) randomised trace estimation, but we found that its estimate variance was too high for a computationally feasible number of random vectors. Thus, we implemented the recent quadratic improvement of this algorithm, which is called Hutch++ (Meyer et al., 2021). Similar to our randomised SVD implementation, this trace estimation algorithm evaluates matrix-vector products of ***A*** with a random vector ***x***, without actually computing ***A***. Instead, an ‘oracle’ function is used (Algorithm 1) that returns the matrix-vector product ***Ax*** by working with the sparse Cholesky factor ***L***^*−*1^, see Algorithm 1. Our implementation of Hutch++ also allows for implicit centring via Algorithm 2.

We have implemented the above algorithms in the open-source `randPedPCA` R package. The package also contains utility methods extending R’s `summary` and `print` functions for pedigree PCA objects, and a function for 3D plots with projections that utilises the optional dependency `rgl`. Additionally, the package contains documentation for all exported functions, it includes three documented example datasets (two of which we describe in the Results section), a vignette, and a test suite to minimise inadvertent changes during development.

## 3 Results

To demonstrate the utility of, and to help building an intuition about, pedigree PCA, we ran `randPedPCA` on simulated data and on a large Labrador Retriever pedigree. We also compared the running time `randPedPCA` to that of a standard PCA implementation.

### 3.1 Simulated data

We generated synthetic pedigree and genotype data for two scenarios (2pop and 4pop) using the forward-in-time simulator, AlphaSimR (Gaynor et al., 2021). For the 2pop scenario, we created two populations, 1 and 2, of 50 individuals each, using coalescent simulation. These populations originated from the same ancestral population 100 generations ago. We saved 11,000 segregating loci across 10 chromosomes. We then added two traits, each with genetic variance of 1.0, and a negative genetic correlation of -0.3 The environmental variance of each trait was set to 2.0 and covariance was set to 0.0. Both traits had 100 causal variants per chromosome. We then selected for trait 1 in population 1 and for trait 2 in population 2, using the top ten individuals for the respective trait, and generating 50 offspring in each population per generation. We maintained this regime for 20 additional generations. In generation 10, we created a hybrid/crossbred population of 50 individuals, using parents from 1 and 2, which had been differentially selected for traits 1 and 2, as described above. Each subsequent generation of the hybrid population had 50 individuals. To generate these, we chose two parents from each, 1 and 2, and six from the hybrid population. From each population, we selected those individuals that scored the highest for a selection index that weighted both traits equally.

In the 4pop scenario, we did not use selection. We started with one panmictic ancestral population of 200 individuals, again generated via coalescent simulation, which we propagated for 10 discrete generations at constant size. After that, we split this ancestral population into four daughter populations of 50 individuals each, without gene flow between them. We then propagated each for nine generations, again at a constant population size, and recorded pedigree and SNP genotypes at neutral markers.

For both scenarios, we ran pedigree PCA with `randPedPCA`. We carried out genotype PCA using R’s built-in `prcomp` function (with centring) on all 11,000 genotypes.

#### 3.1.1 pop scenario

All PCAs of the 2pop scenario (Figure 1) showed a rapid decay of the variance captured by the principal components, with principal component 3 always capturing less than 3% of the total variance. There was a clear difference between PCA on non-centred versus centred data. For the non-centred data, see bottom panels of Figure 1, PC2 captured a comparatively high proportion of the total variance and separated populations 1 and 2. PC2 explains 20% of the variance for the pedigree and 12% for the SNP genotypes. PC1 aligned well with time, that is, generation number of the simulated populations. The respective Spearman correlation coefficients, *ρ*, for PC1 according to the pedigree or the SNP genotypes were 0.88 and -0.90. PC1 captured a large amount of the total variance, 25% in the case of the pedigree PCA and 73% for the genotype PCA.

**Figure 1.**
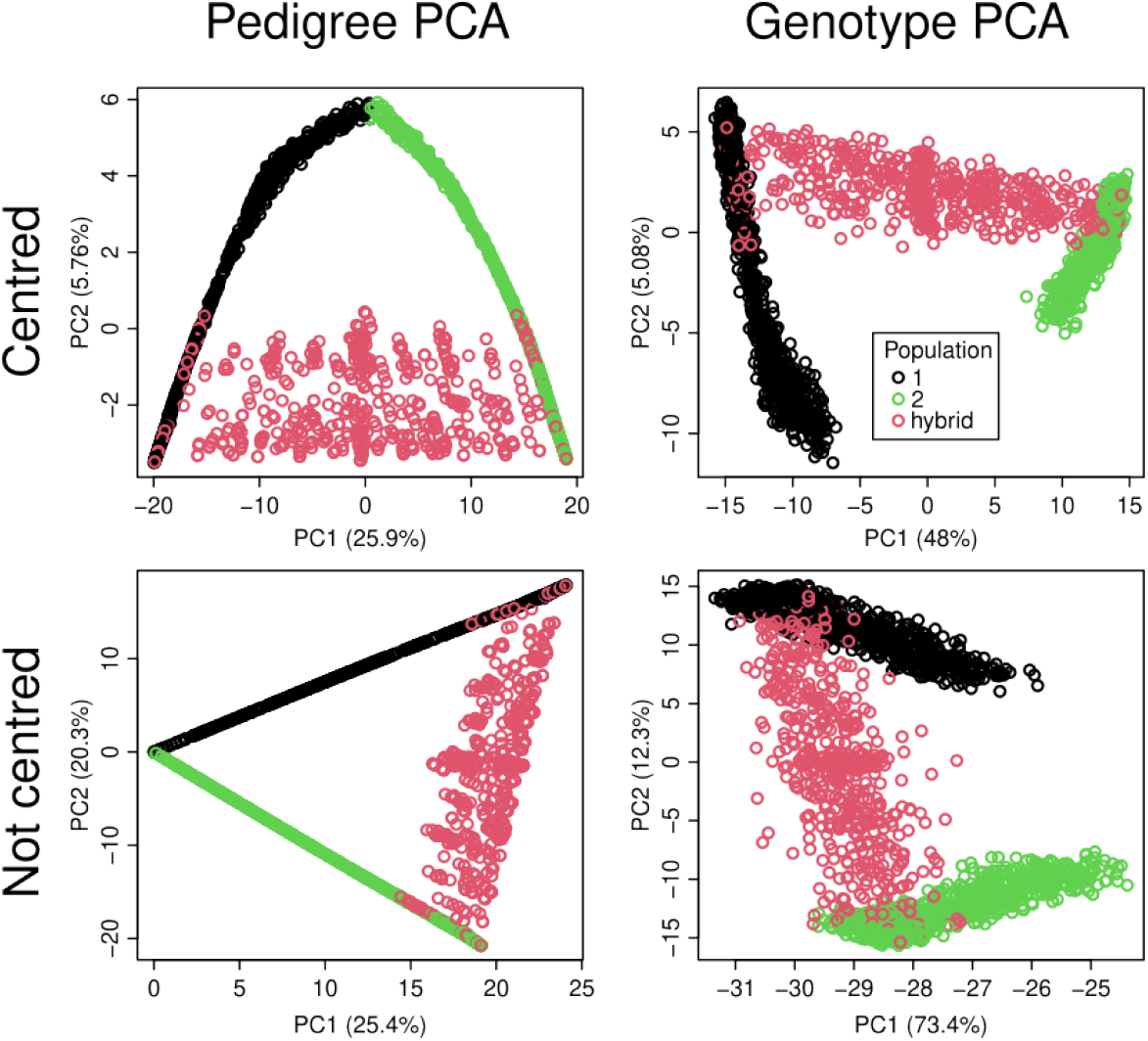
Scatter plots of the first two principal components computed from the pedigree and SNP genotypes of the 2pop scenario with centring (top row) or without centring (bottom row). The plots on the left were generated with `randPedPCA` and thus show approximate scores and percentage of captured variance. We ran the standard PCA on all SNP markers. The legend applies to all panels.

With centring (Figure 1, top panels), there was a greater drop in the variance captured between PC1 and PC2. For the PCAs on the centred data, it was PC1 that separated populations 1 and 2. It captured 26% of the total variance for the pedigree PCA and 48% for the genotype PCA. PC2 captured considerably less of the total variance, 5.7% for the pedigree PCA and 5.1% for the genotype PCA. With centring, it was PC2 that aligned with the generation number, with *ρ* values of -0.97 for the pedigree PCA and 0.86 for the genotype PCA. The top-right panel of Figure 1 shows how the generations of population 1 (in black) align with PC2. The generation of population 2 (in green), also spread out along PC3 (not shown). Comparing the plots of pedigree PCA with those of genotype PCA (the left and right-hand panels of Figure 1), there was a strong resemblance in general shape. The fact that the axes were flipped in both cases owes to the fact that PCA does not preserve the sign of individual principal component score vectors. The most notable difference between plots of the pedigree and the genotype PCA was that the individuals at the top of the pedigree, which are the founders of populations 1 and 2, are shown in the same point in the pedigree PCA plot, despite being genetically diverged. The genotype PCA plots on the right-hand side of Figure 1, however, show populations 1 and 2 as separate.

#### 3.1.2 pop scenario

The PCAs of the 4pop scenario (Figure 2), had variance components that decayed more slowly. Again, without centring (bottom panels), PC1 aligned well with the generation number of the simulated individuals, capturing 6.2% of the total variance for the pedigree and 76% for the SNP genotypes. With centring (top panels), it was the population structure that dominated the top principal components, whereas time aligned only with PC4 (not shown). For both pedigree and SNP genotypes, patterns emerged that resembled a central point cloud (red dots) with four spikes (green, light blue, dark blue, and black dots). The spike ends were arranged like the points of a regular tetrahedron when also taking into account PC3 as the third dimension (not shown). The central cloud contained individuals of the ancestral population, whereas each spike corresponded to one derived population. We observed similar-sized variance components of approximately 4 to 5% for principal components 1 and 2 for both the genotype and the pedigree PCA with centring.

**Figure 2.**
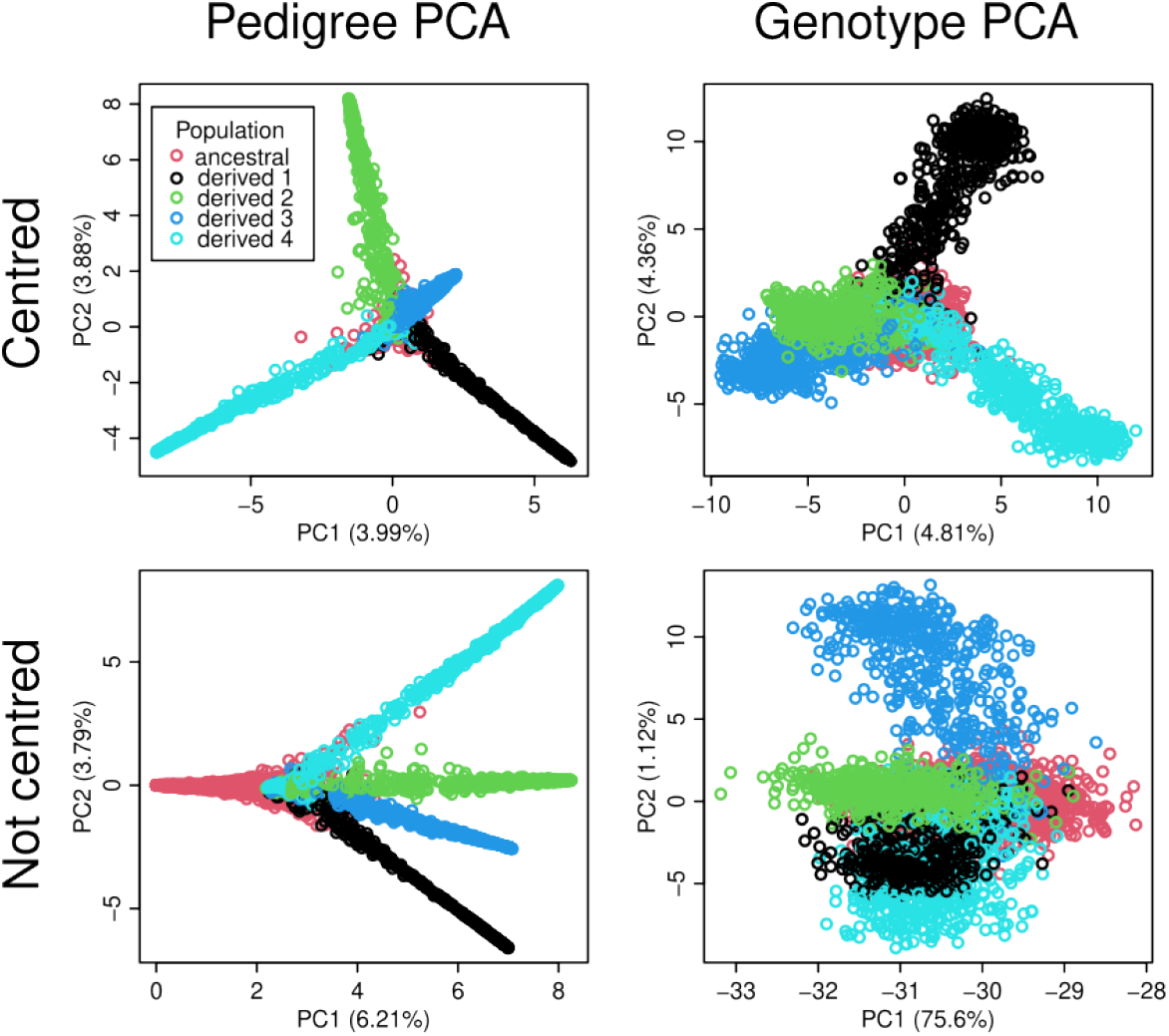
Scatter plots of the first two principal components computed from the pedigree and SNP genotypes of the 4pop scenario with centring (top row) or without centring (bottom row). The plots on the left were generated with `randPedPCA` and thus show approximate scores and percentage of captured variance. We ran the standard PCA on all genotype markers. The legend applies to all panels.

### 3.2 Labrador Retriever data

To demonstrate the application of `randPedPCA` to empirical data, we used the registered canine pedigree of the Labrador Retriever, provided by the Kennel Club, UK. As the most popular breed in the UK, the Labrador Retriever pedigree consists of 1,486,764 records covering 70 years (1955 to 2025). However, because records were not fully digitised until 1990, the pedigree is incomplete, with founders occurring throughout the pedigree. We analysed it as such, without defining unknown parent groups. Data cleaning was minimal, with the addition of individual records for any sires and dams not listed as individuals, and the resolution of two pedigree loops (where an individual is found to be their own ancestor). We identified these loops when sorting the pedigree, and located both offending individuals using the `visPedigree` R package (Luan, 2025). Both pedigree loops had been caused by incorrect sire allocation, which we set to missing. We then reordered the pedigree from the ancestors to the descendants. Generations were counted forward in time, where generation one represents the founders. All non-standard recorded coat colours were set to unknown to compare population structure only for black, yellow, and chocolate coat colours. Finally, we performed a centred pedigree PCA, using `randPedPCA`.

The first three principal components of the Labrador pedigree (Figure 3) captured 6.5% of the total variance, with PC1 capturing 4.1%, PC2 0.70%, and PC3 0.33%. This aligns with a genotype PCA on a representative sample of Kennel Club registered Labradors (Wiener et al., 2017). PC2 was negatively correlated with time, whether measured by generation (*ρ* = *−* 0.63) or year of birth (*ρ* = *−* 0.58), see the right-hand panel of Figure 3, vertical axis. We found an observable grouping of chocolate Labrador Retrievers, shown in brown in the left-hand panel of Figure 3. The finding resonates with other studies that have reported differences between chocolate and non-chocolate (yellow and black) Labradors (e.g. McGreevy et al., 2018; Wiener et al., 2017). Although Labrador Retrievers were recognised as a breed by the Kennel Club in 1903, the chocolate colour was only included in 1930, partly due to the rarity of the chocolate colour. Only recently has their popularity increased. Thus, it is plausible that these show comparatively low scores on PC2, which is negatively correlated with time.

**Figure 3.**
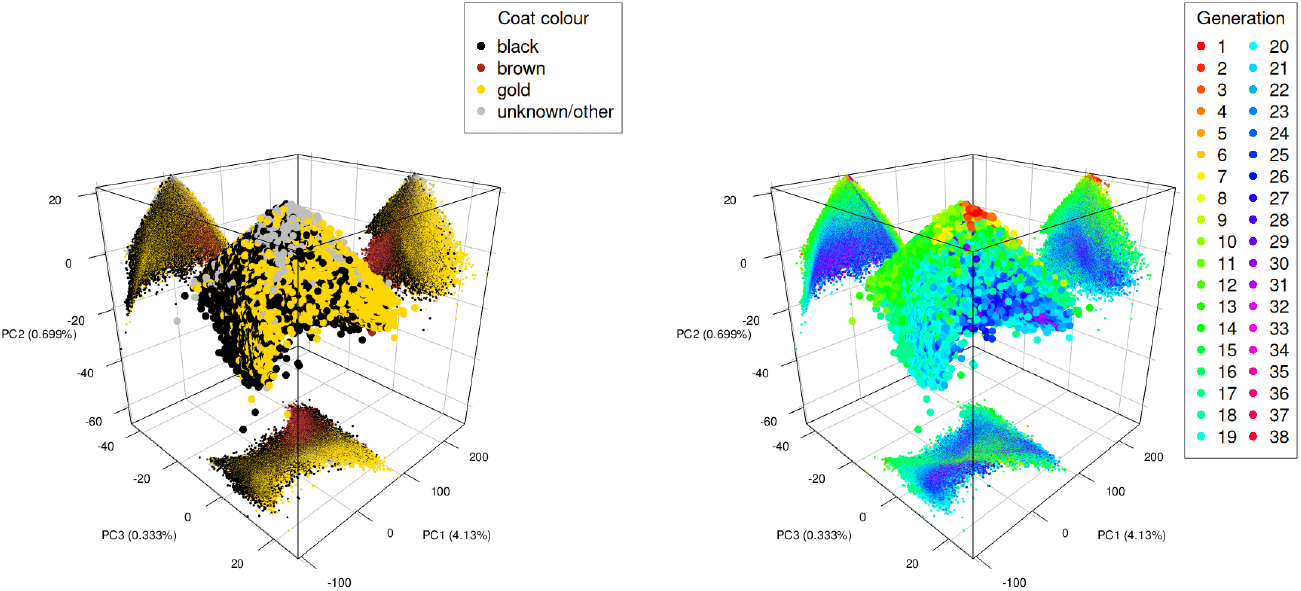
3D scatter plots of the first three principle components for the UK Kennel Club’s Labrador Pedigree from 1955 to 2024. The left-hand plot highlights coat colour, while the right-hand plot highlights generation. Projections onto coordinate planes are provided with the same colour used in the main plots.

### 3.3 Performance

For the simulated 2pop dataset, `randPedPCA` computed an approximate PCA in 8 ms, while R’s naïve `prcomp` took about 42 s, a performance difference of greater than three orders of magnitude (Table 1). We did not attempt to obtain such a running time ratio for the Labrador dataset, as `prcomp` would require the dense ***L*** matrix as input, which itself is prohibitively costly to compute and store from a pedigree with almost 1.5 million individuals. However, our `randPedPCA` finished in under 20 s.

**Table 1:**
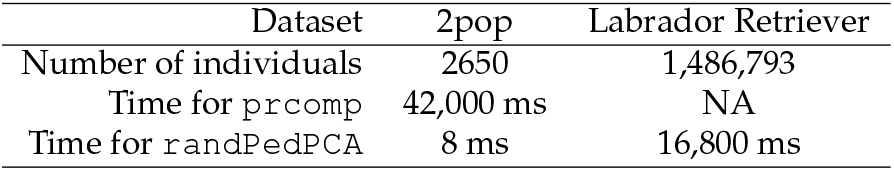
Running times of randPedPCA and R’s built-in prcomp on a standard laptop computer. The datasets are the 2pop simulation and the UK Labrador Retriever pedigree.

| Dataset | 2pop | Labrador Retriever |
| --- | --- | --- |
| Number of individuals | 2650 | 1,486,793 |
| Time for `prcomp` | 42,000 ms | NA |
| Time for `randPedPCA` | 8 ms | 16,800 ms |

## 4 Discussion

In this paper, we introduced a method for rapid PCA of large pedigrees, which complements the existing toolbox for analysing pedigrees (e.g. Aguilar et al., 2011; Colleau, 2002; Garbe and Da, 2003; García-Cortès et al., 2010; Gengler et al., 2007; Gutièrrez and Goyache, 2005; Meuwissen and Luo, 1992; Sargolzaei et al., 2005; Strandèn and Mäntysaari, 2020; Tier, 1999). This is the pedigree counterpart to the popular genotype PCA (Hanotte et al., 2002; Menozzi et al., 1978; Novembre and Stephens, 2008; Patterson et al., 2006). This method is implemented in freely available `randPedPCA` R package. Below, (i) we describe the package’s user interface and performance, (ii) explore the meaning and impact of centring and how pedigree PCA compares to genotype PCA, and (iii) discuss incomplete pedigrees and applications.

### 4.1 User interface and performance

Our R package for fast randomised pedigree PCA and visualisation, `randPedPCA`, is avalable from CRAN. The principal intended use is in Integrated Development Environments such as Rstudio or VSCode. But `randPedPCA` may also be used on the command line or as part of pipelines, for instance, for quality control.

The package’s main user-facing function is called `rppca`. This function can take as input a `pedigreeTools` pedigree object or a sparse ***L***^−1^ matrix in `spam` format. It then performs PCA via an approximate randomised SVD and returns an object of the S3 class `rppca`, which is modelled after R’s built-in `prcomp`. We recommend that `rppca` is used with centring, that is, with the parameter `center` set to `TRUE`.

To inspect the outcome of the PCA, we created the S3 methods `summary.rppca` and `plot.rppca`. Because R’s base plotting is slow with many points, `plot.rppca` down-samples the number of dots to be shown to 10,000. This works via an index vector that is added to the `rppca` object by the function `dspc`. This function may also be run independent from plotting, providing more flexibility as to the number of dots shown and making plots reproducible.

For interactive 3D plotting, we generated a function called `plot3DWithProj`, which depends on the R package `rgl`. Because correctly installing `rgl` can be tedious, we declared it as a suggested package, not a formal dependency of `randPedPCA`. We believe that the 3D plots generated with `plot3DWithProj` are a very valuable data analysis tool as they allow one additional dimension to be concurrently inspected comared to a 2D plot. The common criticism of 3D plots, that it is not possible to know the correct position of a dot, is alleviated by the fact that we show 2D projections of all dots on to the coordinate planes, as shown in Figure 3.

We showed that `randPedPCA` is more than three orders of magnitude faster than a naïve implementation when run on a simulated dataset of moderate size, 2650 individuals (Table 1). PCA on large pedigrees is virtually impossible without randomised linear algebra, and we are not aware of any previous large-scale implementation. Thus, there is no other package to which we can compare `randPedPCA` in a fair way. An alternative to PCA is rendering a graph based on a pedigree’s relationship matrix, ***A***, or the adjacency matrix, ***T*** ^*−*1^. This is also considerably slower than generating a PCA plot with `randPedPCA` (Table 1). Our work is related to Garbe and Da (2003) who visualised large pedigree graphs by collapsing large progeny groups into nodes. Their approach retains the original display of a pedigree, while our approach projects pedigree information into lower dimensions. Our work is also related to Feng et al. (2018) who developed a fast randomised PCA for sparse data and Erichson et al. (2019) who developed the `rsvd` R package for general matrices, but we leverage the well-known result about the sparsity of the pedigree precision matrix and its Cholesky factor for a scaleable PCA of large pedigrees. Lastly, our work is also related to scaling PCA to large genomic datasets, which are increasingly available in breeding programmes and biobanks. To this end, Galinsky et al. (2016) and Li et al. (2023), and Miles et al. (2024) implemented randomised SVD, while Agrawal et al. (2020) implemented probabilistic PCA.

### 4.2 What is the data matrix in pedigree PCA and should it be centred?

PCA is usually performed on a data matrix that has features in columns and individuals in rows. A good example is Anderson’s popular iris dataset, which comprises measurements of four flower traits taken from 150 individuals of three species of iris (Fisher, 1936). It is often advised to centre the data matrix before PCA (e.g. Jolliffe, 2002). That is, the column means should be made equal to 0. One reason for this is that, when performing PCA on a dataset with non-zero column means, the first principal component tends to represent all column means’ differences from zero. PCA is usually carried out via (1) eigen decomposition of a covariance matrix of the features or (2) SVD of the data matrix. Alternatively, PCA may also be carried out via (3) eigen decomposition of the covariance matrix of the individuals. This third path is often not useful for ‘tall’ datasets with more individuals than features, like the iris data, because it involves the decomposition of the large covariance matrix of individuals rather than the smaller covariance matrix of the features. It is useful, however, for feature-rich ‘wide’ datasets such as omic data where there are often many more features than individuals (e.g. Pocrnic et al., 2022). There is another key point where approach (3) differs from (1) and (2). The approaches (1) and (2) create, among other things, a matrix of loadings, called the eigen vectors or right singular vectors, depending on the method. This loadings matrix has then to be multiplied with the original data matrix to obtain the principal component scores. With approach (3), however, there is no need to go back to the data matrix. Rather, if we decompose the covariance matrix of individuals, the principal component scores are equal to the matrix of the left singular vectors, ***U***, multiplied by the diagonal matrix of the squared singular values, diag(*d*^2^). This is also the approach we used when generating a pedigree PCA, based on implicitly decomposing the additive relationship matrix ***A***, which is a covariance matrix of individuals.

Following the standard advice, genotype PCAs are usually based on centred allele dosages, although other implied statistical models and corresponding covariance matrices are also possible (e.g. Legarra, Bermann, et al., 2024b; Powell et al., 2010). Unlike for genomic data with individuals in rows and loci in columns, it is not immediately clear what is the corresponding data matrix for a pedigree and whether one should centre it for PCA. One can encode a pedigree graph as an adjacency matrix, though this representation does not imply a statistical model with a corresponding covariance matrix.

For genomic data, the relationship matrix is the covariance matrix, 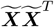, of the (centred) genotype data, 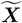. Hence, working back from the pedigree relationship matrix, a data matrix ***D*** should fulfil the equation ***DD***^*T*^ = ***A***. Many matrices fulfil this criterion, one of them being ***L***, the Cholesky factor of ***A***. The matrix ***L*** encodes the relations of each individual with all of its ancestors, following the pedigree quantitative genetic model (Kennedy et al., 1988; Quaas, 1988; Thompson, 1979; Wright, 1921, 1922). This model treats the pedigree founders as a ‘reference’ (base) population with additive genetic values distributed with mean zero and base population additive genetic variance. The non-founder values are then recursively modelled as a deviation from the average value of their parents, in line with recombination and segregation of parental genomes. Hence, the pedigree founders serve as the ‘centring’ point. However, the matrix ***L*** could be centred differently for PCA, depending on the aims of the analysis. While it is computationally costly to compute and store ***L*** for large pedigrees, we have developed a scalable algorithm for such centring, which can also be used within pedigree PCA (Algorithm 2).

We found that plots of principal component scores obtained with and without centring commonly had swapped principal components 1 and 2. Without centring, principal component 1 tended to align with time or generation number, while accounting for a large amount of variance. With centring, time tended to covary with a lower-order principal component and population structure was reflected in the leading principal component(s). Centring also made plots of pedigree PCA look more alike those of genotype PCA, which are generally computed on a centred genotype matrix. Whether we centre or not and which individuals we chose as the ‘reference’ population, is related to our choice of the reference/mean point from which covariance is defined and genetic relationships interpreted (Legarra, 2016; Legarra, Bermann, et al., 2024b; Legarra, Christensen, et al., 2015; Powell et al., 2010). For example, Fan et al. (2022) has recently used the realised identity-by-descent information from an ancestral recombination graph (pedigree gives expected identity-by-descent) and performed PCA with different time-depths to study how population structure changed over time. Our implementation centres across all individuals, but Algorithm 2 can be changed to use just a group of individuals, which we leave for future work.

### 4.3 Real world pedigrees and applications

One notable difference between the genotype and pedigree PCA plots was that for pedigree PCA all founding individuals were placed at the same location. This was obvious for the simulated 2pop scenario and the Labrador Retriever example. For both, the scores of the leading principal components were essentially identical for the founding individuals. This is plausible given the logic that PCA summarises common patterns in higher-order principal components. Pedigrees provide information about the expected provenance of the pedigreed individuals’ genetic material. The further ‘away’ from the founders an individual is located, the more information there is about that individual and its relationships to other individuals. However, there is no such information for individuals with no known parents, the founders of a pedigree. In the relationship matrix, the covariances between the founders are zero. Thus, for the sake of PCA, these individuals lack shared patterns that could be summarised. These individuals then contribute low-order principal components, each to their own. But only the leading components are plotted as only those contain covariance patterns shared across many individuals.

This behaviour is not necessarily an issue, in particular when there are few founders which occur only in early generations. One straightforward fix would be to exclude any founder individuals from the PCA plot. If there are any external data on or assumptions about the genetic differentiation between founders (Kennedy et al., 1988; Legarra, Bermann, et al., 2024a; Macedo et al., 2022; Wicki et al., 2023), the concept of metafounders (Legarra, Bermann, et al., 2024b; Legarra, Christensen, et al., 2015) may be used to group founders according to this information. For example, a pedigree with founders belonging to two (sub-)populations may be extended with two metafounders with appropriate covariance between them (Legarra, Christensen, et al., 2015). Pedigree PCA would then differentiate the founders between the two (sub-)populations. In this case, the metafounders, and not the founders, would end up with very similar scores in their leading principal components. Again, they could be omitted from the plot.

In addition to the visualisation of large pedigrees, our work will complement existing approaches to define and study sub-populations in breeding programmes (e.g. Anglhuber et al., 2024; Steyn, Lawlor, et al., 2023; Steyn, Masuda, et al., 2022). It will also complement approaches to optimise selection of key individuals in conservation, genotyping, genome sequencing, and large-scale genomic estimation of breeding values (e.g., see Pocrnic et al., 2022, and references therein). The logical extension of the approach we presented here is to combine pedigree and genotype data using the ‘single-step’ covariance matrix (Legarra, Aguilar, et al., 2009). We have made initial progress towards this, leveraging the work of Colleau et al. (2017). The full application is beyond the scope of the present paper.

## 5 Conclusions

Visualising large pedigrees is a long-standing and challenging problem. Here we introduce the `randPedPCA` R package that rapidly performs PCA on large pedigrees. This package thereby enables a straightforward and scaleable visualisation of large pedigrees. When such a PCA is combined with metadata, we can clearly study the structure of large pedigrees and highlight their key drivers of variation. The `randPedPCA` R package is freely available from CRAN and GitHub. In addition to the visualisation of large pedigrees, our work will complement existing approaches to define and study sub-populations in breeding programmes. It will also complement approaches to optimise selection of key individuals in conservation, genotyping, genome sequencing, and large-scale genomic estimation of breeding values.

## 6 Declarations

### 6.1 Ethics approval and consent to participate

Does not apply.

### 6.2 Consent for publication

Does not apply.

### 6.3 Availability of data and material

The `randPedPCA` R package is available on CRAN TODO (stable/published version) and GitHub https://github.com/HighlanderLab/RandPedPCA (stable and development versions). The R code for simulation of data and demonstration of the package is available on GitHub in the directory `data-raw` of the R project folder. The anonymised Labrador Retriever pedigree was provided by The Kennel Club. `randPedPCA` R package depends on the `spam`, `pedigreeTools`, and `Matrix` R packages. The `rgl` R package is required to make use of the 3D plotting function, but it is not a formal dependency.

### 6.6 Authors’ contributions

HL derived the algorithm, wrote a Python prototype, and contributed to writing the manuscript. RFC ran the real data example and provided feedback on the manuscript. GG led the project, initiated the simulation, and contributed to the writing of the manuscript. HB developed the `randPedPCA` R package, extended the simulations, and wrote the initial draft of the manuscript. All authors have read the final version of the manuscript.

## 6.7 Acknowledgements

HL acknowledges support from Departmental Fellowship at the University of Michigan, Department of Statistics, Ann Arbor. RFC acknowledges support from the The Kennel Club and the University of Edinburgh. We are grateful to Joanna Ilska, Daniel Tolhurst, Ivan Pocrnić, Jarrod Hadfield, and Andres Legarra for their helpful input to this project.

